# The Effect of Plosive Content on the Loudness Perception of Vowel-Consonant-Vowel Syllables in Listeners with Sensorineural Hearing Loss

**DOI:** 10.64898/2026.08.26.747282

**Authors:** Thomas Davis, Stefan Bleeck

## Abstract

**Objective:** This study investigated whether plosive consonants carry a perceptual loudness weighting that significantly exceeds that of non-plosive consonants when judged by hearing-impaired listeners.

**Design:** A prospective loudness matching experiment utilizing the method of adjustment.

**Study Sample:** 19 consenting native English speakers (Mean age: 61.4, SD: 16.4) with bilateral mild to moderate high-frequency sensorineural hearing loss, indicative of presbycusis.

**Stimuli:** 13 vowel-consonant-vowel (VCV) nonsense syllables, exclusively utilizing the flanking vowel /u/.

**Results:** Descriptive analysis revealed a strong time-order effect influencing loudness judgments for 7 of the 13 VCV test stimuli. Statistical testing showed no significant difference (*P* = 0.94) between the relative amplitudes corresponding to the point of equal loudness for plosive-containing versus non-plosive-containing VCV stimuli. However, 6 individual VCV stimuli, containing consonants from 4 separate manners of articulation, produced significant loudness matching data (*P <* 0.01).

**Conclusions:** The results falsify the hypothesis that plosives, analyzed collectively as a class, possess a heavier perceptual loudness weighting than non-plosive consonants. While 6 individual VCV stimuli indicated potential individual consonantal loudness weightings, these findings must be interpreted cautiously due to the restriction to a single vowel context and the presence of procedural time-order biases.

## 1 Introduction

The speech banana (Lidén and Fant, 1954; Fant, 2004) is a well-established clinical and educational heuristic that maps the intensities and representative frequencies of speech sounds—which collectively constitute the long-term average speech spectrum (LTASS)—onto an audiogram referenced in hearing level (dB HL). To construct conventional speech bananas, representative phonemic spectral peaks are identified and converted from sound pressure levels (dB SPL) to dB HL using standard Reference Equivalent Threshold Sound Pressure Level (RETSPL) values (ANSI, 1996; Yost and Killion, 1997). By comparing these phonemic levels against pure-tone thresholds, clinicians obtain a visual estimate of whether individual speech sounds are likely to be audible during conversational speech (Boothroyd et al., 1994).

However, representing running speech purely through a free-field LTASS carries recognized acoustic and psychophysical limitations. Free-field to eardrum transfer functions and outer/middle ear resonances naturally shape the spectral envelope, effectively flattening the functional representation of speech at the tympanic membrane (Pascoe, 1980; Hawkins, 1984; Amos and Humes, 2007). Furthermore, running speech contains dynamic short-term spectral characteristics that are obscured by long-term spectral averaging (Turner, 1993). While the conventional +12 dB dynamic range allowance above the LTASS RMS level serves as a standard approximation for Articulation Index calculations (Dunn and White, 1940; French and Steinberg, 1947; ANSI, 1997), instantaneous peak-to-RMS ratios of isolated consonants can exceed these values substantially (Stelmachowicz et al., 1993; Boothroyd et al., 1994).

Beyond physical acoustics, the psychophysical perception of speech loudness is characterized by complex loudness dominance and integration across spectral components (Berg, 1990; Oberfeld et al., 2013). A fundamental question is whether the perceived loudness of a syllable is governed almost exclusively by high-energy vowel formants or whether consonantal segments contribute a distinct perceptual weighting (Lehiste and Peterson, 1959; Montgomery et al., 1987). While earlier studies suggested that speech loudness perception is predominantly vowel-dominated (Montgomery et al., 1987), subsequent work demonstrates that altering consonant-to-vowel intensity ratios can systematically shift overall loudness judgments (Orr et al., 2010). Consonants encompass diverse acoustic classes—including plosives (stops), fricatives, nasals, liquids, and semi-vowels—each exhibiting distinct temporal envelopes and crest factors (Santini et al., 2016). Among these, plosives (/p/, /t/, /k/, /b/, /d/, /g/) possess unique transient burst characteristics and silent intervals. In an early rating study using consonant-vowel (CV) syllables, Sharf (1971) reported that CV syllables containing plosives received higher average loudness ratings than syllables containing other consonant classes. However, whether this effect reflects a general perceptual loudness weighting that persists across different phonetic contexts remains an open question.

Most previous investigations evaluated consonants in syllable-initial (CV) positions or in meaningful words (Sharf, 1971; Orr et al., 2010). In running speech, however, consonants frequently occur in intervocalic contexts. Investigating vowel-consonant-vowel (VCV) nonsense syllables allows the isolated evaluation of consonantal loudness contributions while controlling for lexical familiarity and vowel boundary energy. Furthermore, because individuals with sensorineural hearing loss exhibit altered loudness growth and reduced spectral resolution, evaluating phonemic loudness weightings in hearing-impaired listeners provides insight into how phoneme-level intensities are processed in impaired auditory systems.

The present exploratory study investigates whether plosives carry a distinct perceptual loudness weighting relative to non-plosive consonants in symmetrical VCV syllables presented to hearing-impaired adult listeners. By systematically adjusting consonant level within fixed-vowel VCV stimuli using the method of adjustment, we test whether plosive content systematically biases overall syllable loudness matching.

## 2 Methods

### 2.1 Participants

Nineteen native English-speaking adults (10 females, 9 males; mean age = 61.4 years, SD = 16.4) participated in the main experiment. Pure-tone audiometry (PTA) was conducted in an audiometric sound booth across standard octave frequencies (0.25–8 kHz). All participants presented with bilateral, mild-to-moderate high-frequency sensorineural hearing loss typical of presbycusis, with symmetrical air-conduction thresholds and no air-bone gaps exceeding 10 dB HL. Participant demographic details and mean bilateral pure-tone thresholds are summarized in Table 1 and Table 2.

**Table 1:**
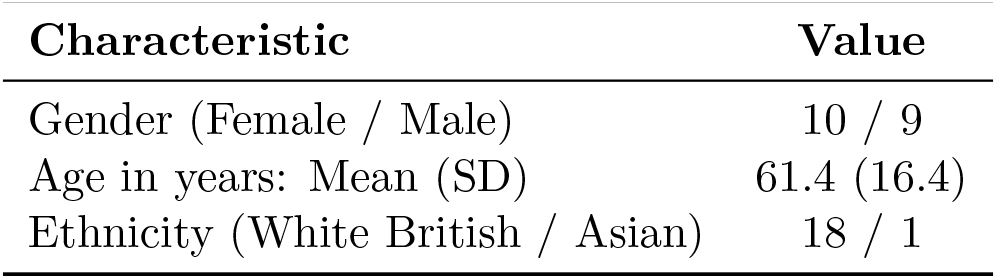
Participant demographic characteristics (*n* = 19).

**Table 2:** Mean pure-tone air-conduction hearing thresholds (dB HL) across audiometric frequencies for the hearing-impaired cohort (*n* = 19).

| Frequency (Hz) | Right Ear Mean (SD) | Left Ear Mean (SD) |
| --- | --- | --- |
| 250 | 15.2 (12.5) | 16.5 (8.6) |
| 500 | 26.3 (12.7) | 26.3 (10.3) |
| 1000 | 33.4 (17.2) | 32.3 (15.6) |
| 2000 | 39.2 (22.3) | 40.5 (19.0) |
| 3000 | 43.6 (20.2) | 46.8 (15.2) |
| 4000 | 55.7 (19.3) | 50.0 (21.2) |
| 6000 | 52.8 (20.4) | 52.6 (20.9) |
| 8000 | 62.1 (20.7) | 58.4 (21.0) |

Inclusion criteria required: (1) native English proficiency to control for language-dependent phonemic weighting; (2) symmetrical high-frequency sensorineural hearing loss; (3) sufficient unaided speech audibility at conversational levels (65 dB SPL); and (4) absence of bothersome tinnitus, hyperacusis, or recent conductive middle-ear pathology. Prior to experimental testing, otoscopy was performed by a certified clinical audiologist to verify clear external auditory canals. Ethical approval was granted by the Faculty of Engineering and the Environment Ethics Committee at the University of Southampton, and written informed consent was obtained from all participants.

### 2.2 Stimuli and Signal Preparation

The speech material comprised 14 vowel-consonant-vowel (VCV) nonsense syllables recorded by a single adult male native English speaker with a standard general accent. Symmetrical VCV syllables were selected to evaluate intervocalic consonants in isolation while minimizing lexical and semantic biases. The close back rounded vowel /u/ was utilized across all tokens.

The semi-vowel syllable /uju/ served as the fixed reference stimulus across all trials. Thirteen VCV test stimuli were generated by pairing /u/ with consonants spanning four manners of articulation and representative spectral regions:

- **Plosives:** /b/, /d/, /g/, /p/, /t/, /k/
- **Fricatives:** /f/, /s/, /S/, /T/, /z/
- **Nasals:** /m/
- **Liquids:** /l/

Acoustic segmentation and signal manipulation were performed in MATLAB (MathWorks, Natick, MA, USA). Phoneme boundaries were determined via combined visual inspection of the wideband spectrogram and waveform zero-crossings. To isolate the consonant as the sole acoustic variable influencing perceived syllable loudness, the root-mean-square (RMS) level of the flanking /u/ vowels was normalized across all reference and test stimuli. Temporal durations for each vowel and consonant segment were cataloged (Table 3). Consonant amplitude was varied parametrically in 2 dB increments spanning *−*8 dB to +8 dB relative to nominal level, utilizing raised-cosine ramps at phoneme boundaries to eliminate spectral splatter or audible clicks during level adjustments.

**Table 3:**
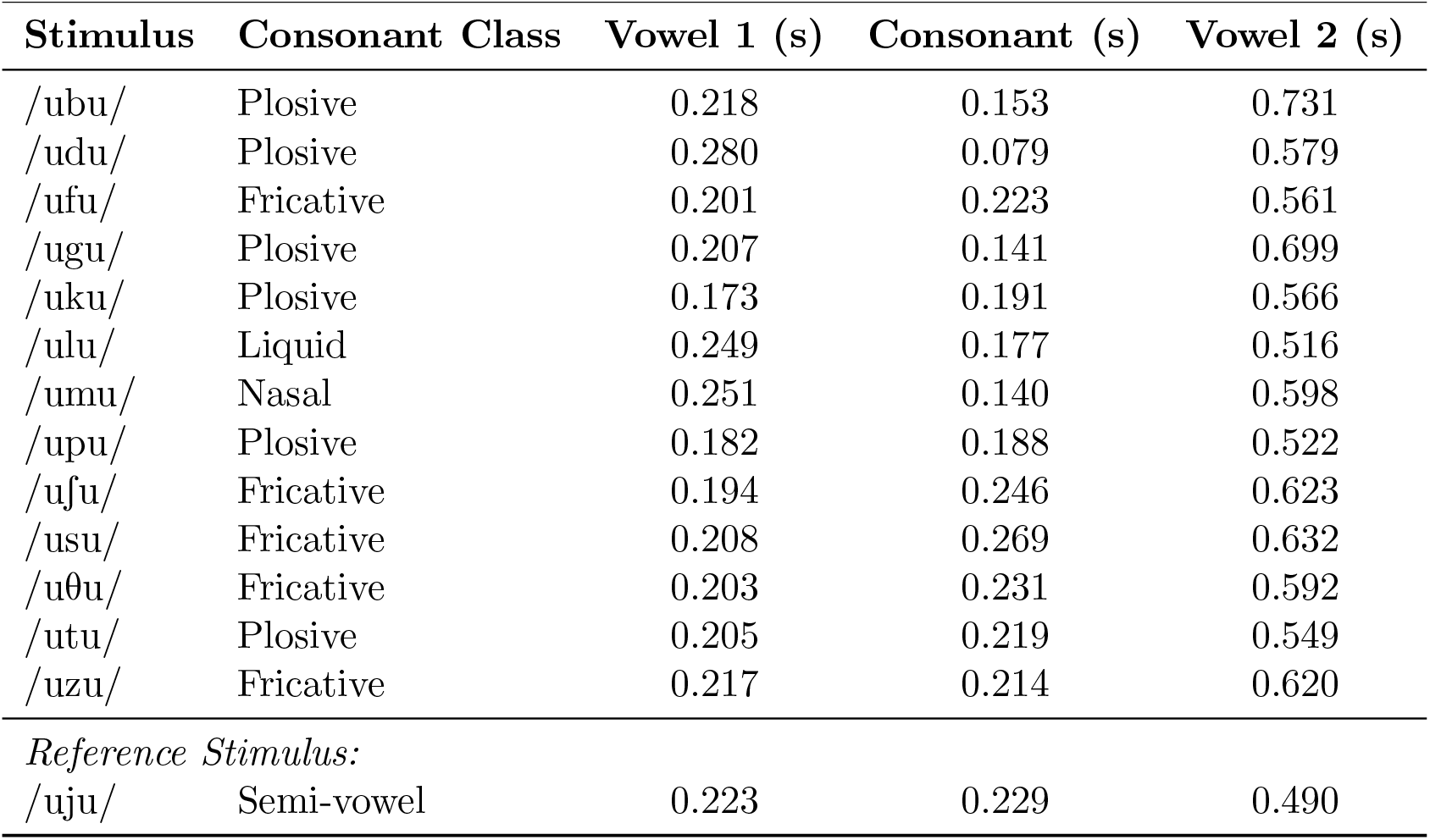
Acoustic segmentation and durations of VCV test and reference stimuli.

### 2.3 Apparatus and Calibration

Acoustic presentations were conducted in a dedicated, sound-treated project room at the Institute of Sound and Vibration Research (ISVR). The participant was seated 80 cm directly in front of an active loudspeaker (Sandstrøm 2.0 Multimedia) at a 0^*°*^ azimuth and ear-level height (119 cm), as illustrated in ETSB Figure I. Experimental sequencing and response acquisition were managed using custom software on a Microsoft Surface Pro tablet.

**Figure 1.**
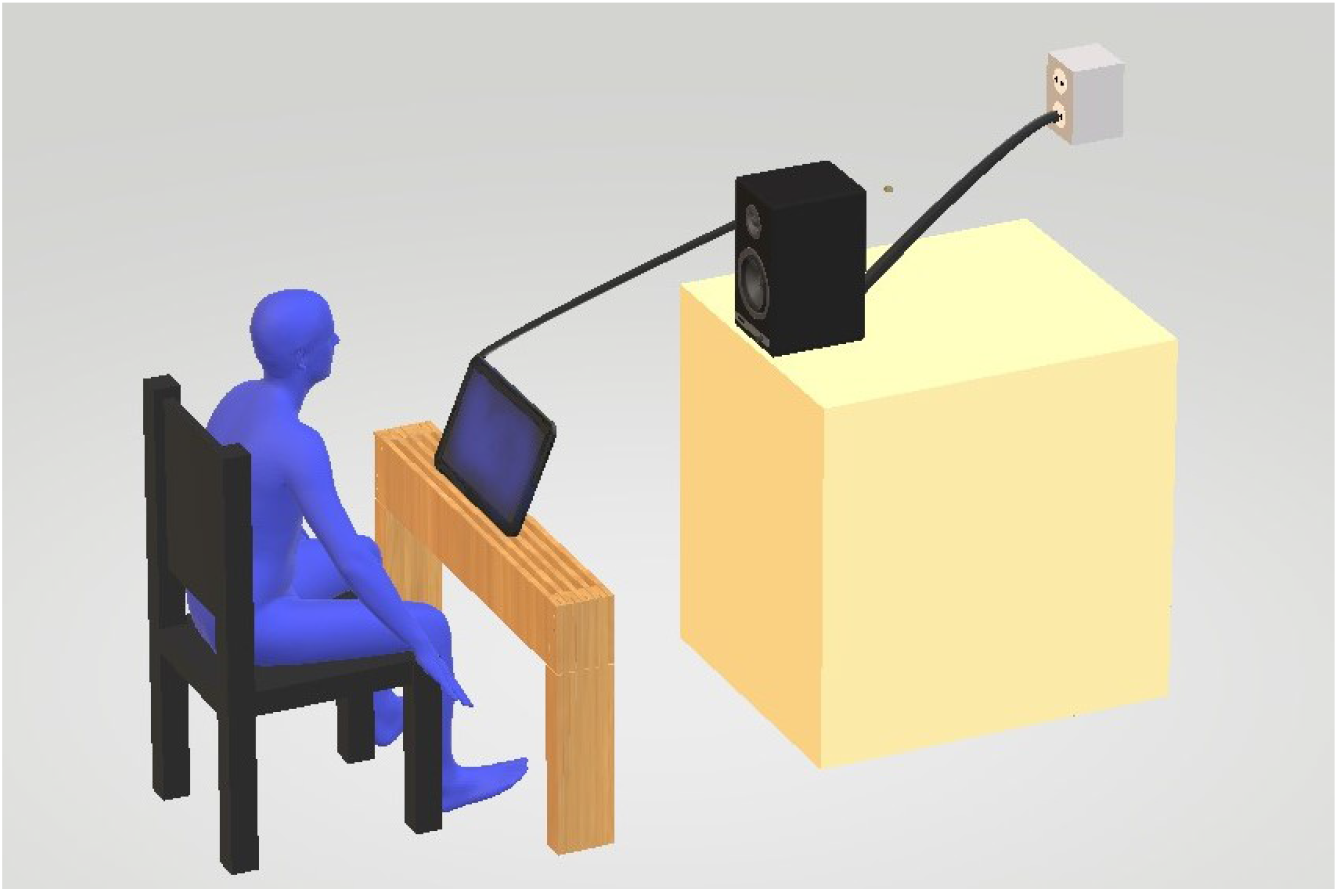
Virtual recreation of the ETSB VCV loudness matching experiment (colour online).

Sound field calibration was conducted using a Class 1 Sound Level Meter (Bruel & Kjaer Type 2250-S) fitted with a 1/2-inch free-field microphone (Type 4189), positioned at the listener head location. The ambient room noise floor produced an equivalent continuous C-weighted level (*L*_Ceq_) of 45.8 dB SPL. Calibration was performed using a 60 s concatenated loop of all 14 VCV stimuli, adjusted to an overall output level of 65.2 dB SPL (*L*_Ceq_), providing a nominal experimental signal-to-noise ratio (SNR) of 19.4 dB.

### 2.4 Loudness Balancing Procedure

A prospective loudness matching paradigm using the method of adjustment was implemented. Each experimental trial presented a paired sequence consisting of the reference syllable /uju/ and a test VCV syllable, separated by a 500 ms inter-stimulus silent interval (ETSB Figure II).

**Figure 2.**
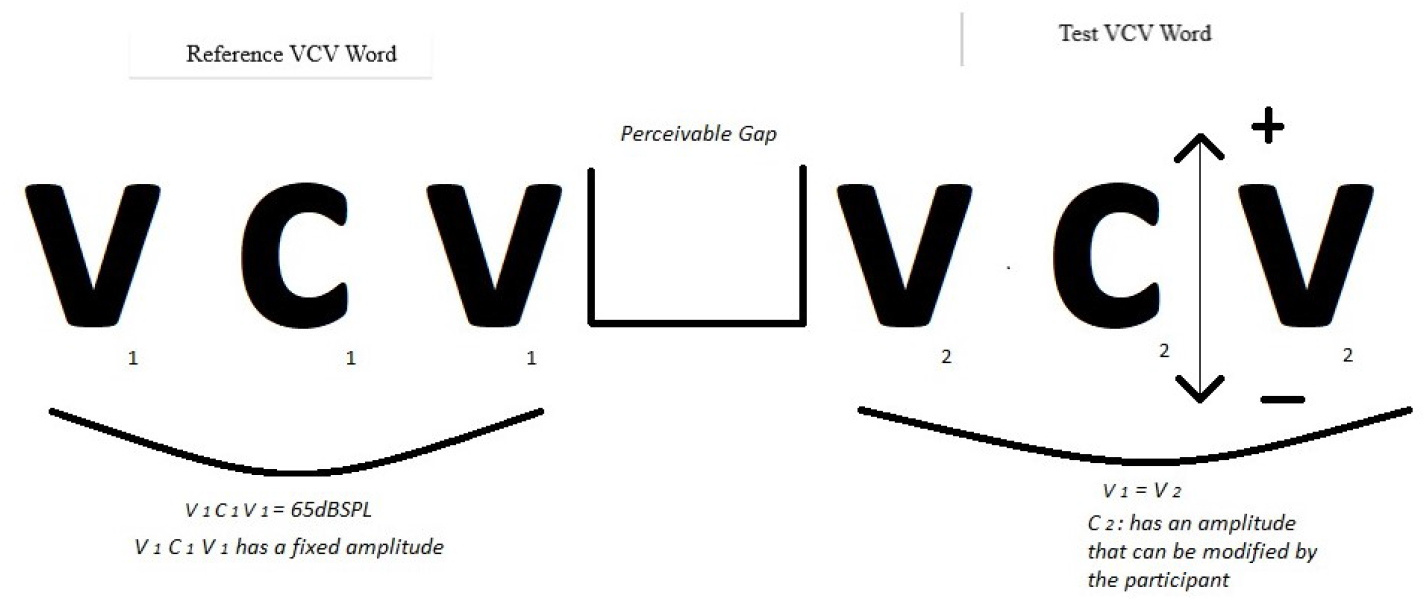
Visual representation of a typical configuration for the VCV word pairs used in the ETSB loudness matching experiment.

To measure and control for time-order biases, presentation order was fully counterbalanced:

1. **Reference-first (Forward):** Reference /uju/ presented in Interval 1; test VCV presented in Interval 2.
2. **Test-first (Reverse):** Test VCV presented in Interval 1; reference /uju/ presented in Interval 2.

Participants adjusted the loudness of the variable interval using a 9-button touchscreen graphical user interface (GUI). The interface displayed adjustment steps corresponding to *−*8, *−*6, *−*4, *−*2, 0, +2, +4, +6, and +8 dB relative to nominal consonant amplitude (ETSB Figure III). To prevent visual bias, numerical decibel values were hidden from participants during testing, and the interface opened at the central neutral position (0 dB) at the onset of each trial. Listeners were instructed to audition alternative levels freely until the overall loudness of the two intervals matched, before confirming their selection.

**Figure 3.**
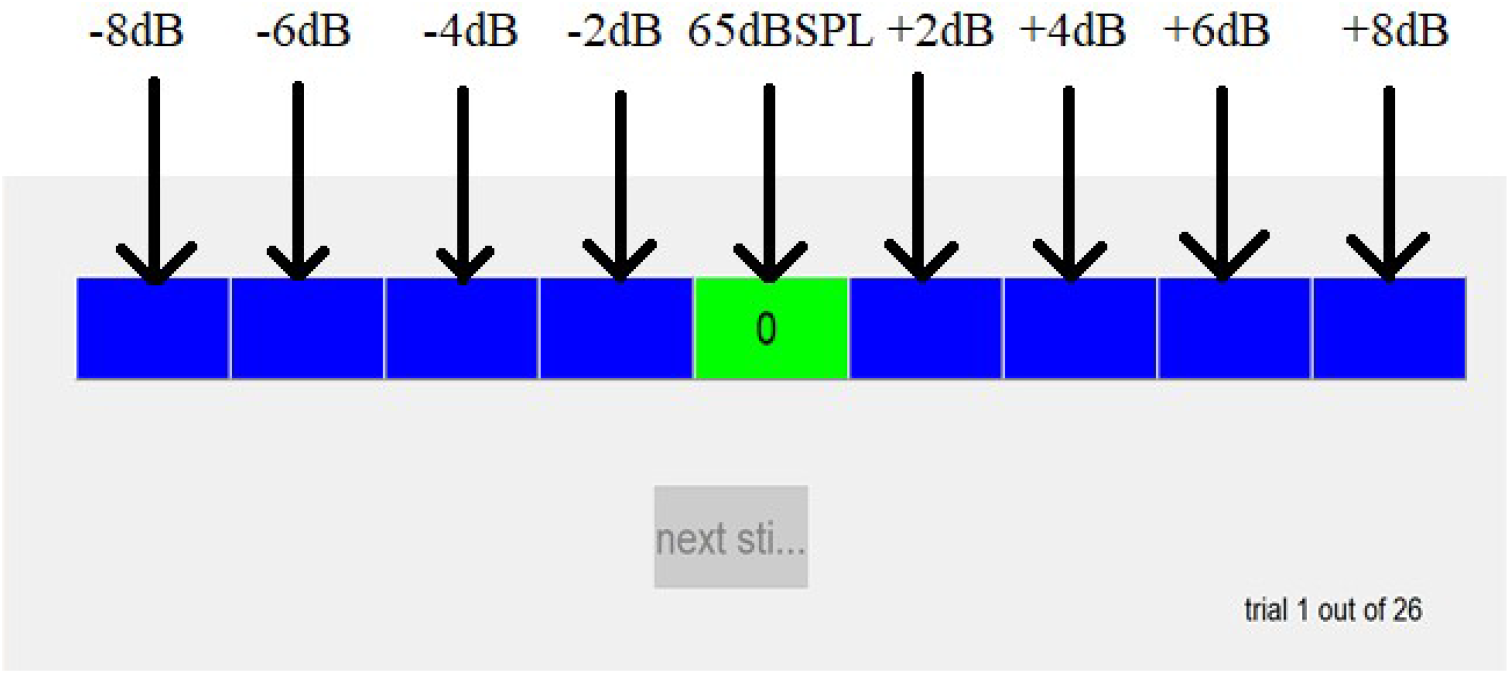
Screenshot showing the ETSB GUI, with arrows pointing to the corresponding physical sound pressure levels of the VCV words; which were produced when the participants pushed each button (colour online).

Prior to testing, participants completed a familiarization run using the continuous calibration signal to confirm comfortable audibility and interface proficiency. Each experimental run comprised 26 randomized stimulus pairs (13 test consonants *×* 2 presentation orders). Participants completed 3 repeated runs, yielding a total of 78 loudness matches per subject (1,482 judgments across the main experimental cohort).

### 2.5 Normal-Hearing Pilot Experiment

To establish baseline psychophysical performance and quantify procedural time-order errors, an independent pilot group of 8 normal-hearing young adults (bilateral thresholds *≤* 20 dB HL across 0.25–8 kHz) completed the identical 78-trial matching protocol. The pilot data confirmed robust execution of the method of adjustment but demonstrated systematic time-order shifts, wherein the second stimulus interval was consistently perceived as louder, necessitating formal integration of presentation order as a fixed within-subject factor in the main statistical model.

## 3 Results

### 3.1 Omnibus Analysis: Plosive versus Non-Plosive Syllables

To test the primary hypothesis that plosive consonants possess an inherent perceptual loudness weighting exceeding that of non-plosive consonants, equal-loudness adjustment levels were first aggregated across consonant classes and presentation orders. The resulting relative adjustment levels (in dB relative to the nominal 65 dB SPL reference) across the four primary conditions are illustrated in ETSB Figure IV.

**Figure 4.**
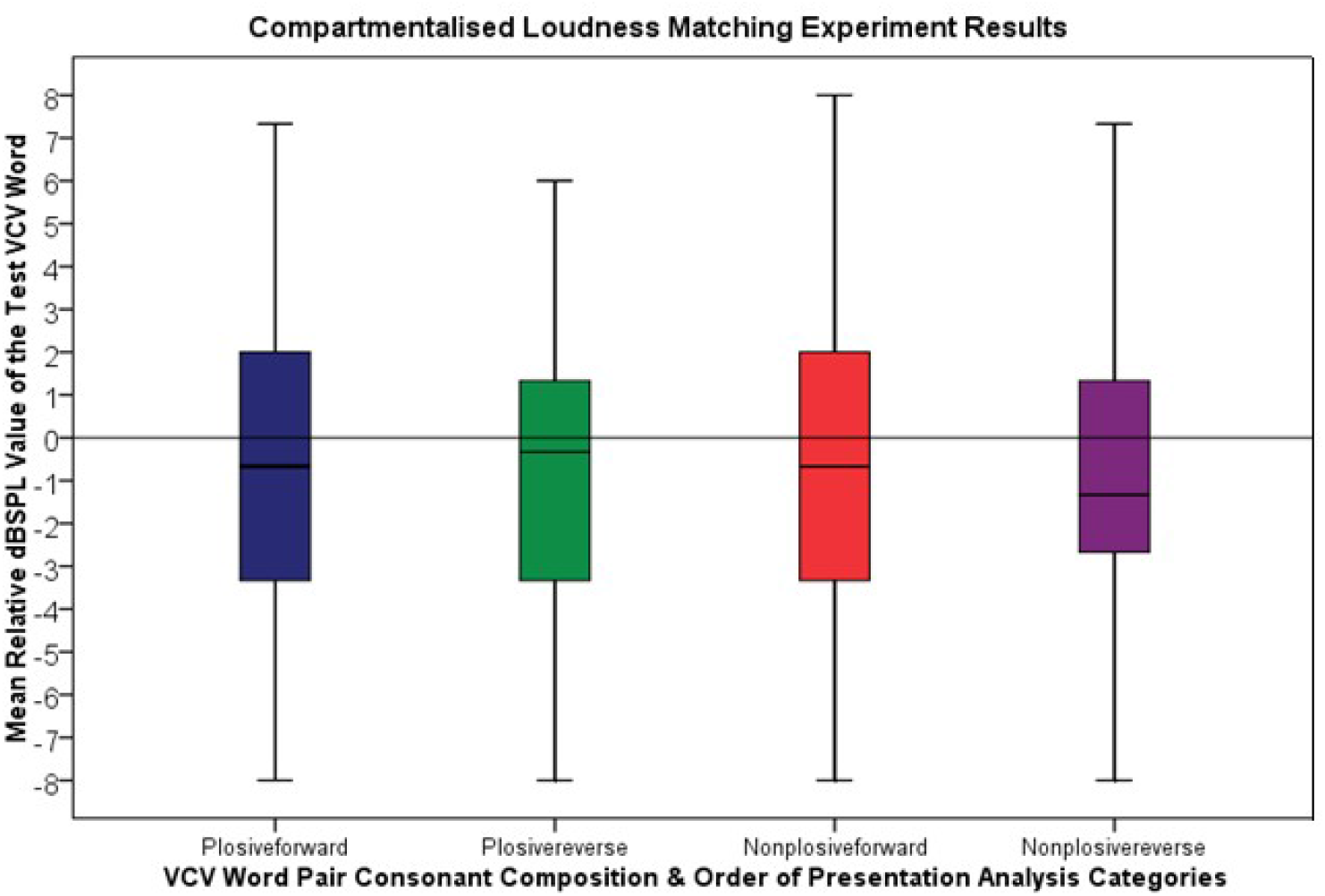
Box and Whisker Plots representing the results of the hearing-impaired participants loudness matching experiment, generalised across four different testing arrangements.

Normality of the distribution of adjustment values across conditions was assessed using the Shapiro-Wilk test (Table 4). With the exception of a minor deviation in the Plosive-reverse condition (*P* = 0.04), the distributions showed no significant departure from normality (*P >* 0.05). Given the established robustness of repeated-measures Analysis of Variance (ANOVA) to moderate departures from normality (Blanca et al., 2017), parametric inferential testing was conducted.

**Table 4:** Shapiro-Wilk test of normality across experimental conditions (*n* = 19).

| Condition | Statistic ($W$) | Significance ($P$) |
| --- | --- | --- |
| Plosive (Forward Order) | 0.932 | 0.16 |
| Plosive (Reverse Order) | 0.898 | 0.04 |
| Non-Plosive (Forward Order) | 0.947 | 0.28 |
| Non-Plosive (Reverse Order) | 0.960 | 0.47 |

A 2*×*2 repeated-measures ANOVA was performed with within-subject factors of **Consonant Class** (Plosive vs. Non-Plosive) and **Presentation Order** (Forward vs. Reverse). As detailed in Table 5, there was no significant main effect of Consonant Class 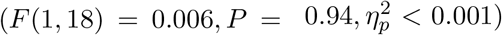. Syllables containing plosive consonants were adjusted to approximately the same relative sound pressure level as syllables containing non-plosive consonants to achieve equal loudness with the /uju/ reference.

**Table 5:** Two-way repeated-measures ANOVA evaluating the effects of Consonant Class and Presentation Order on equal-loudness adjustment levels.

| Source | Type III SS | df | Mean Square | $F$ | Significance ($P$) |
| --- | --- | --- | --- | --- | --- |
| Consonant Class | 0.038 | 1, 18 | 0.038 | 0.006 | 0.94 |
| Presentation Order | 0.925 | 1, 18 | 0.925 | 0.117 | 0.73 |
| Consonant Class × Order | 2.115 | 1, 18 | 2.115 | 0.917 | 0.34 |

Furthermore, the main effect of Presentation Order was not statistically significant 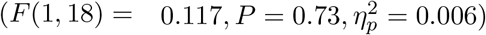, and there was no significant interaction between Consonant Class and Presentation Order 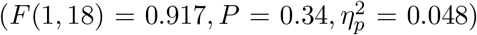. These findings fail to reject the null hypothesis and demonstrate that plosives, evaluated collectively as a phonetic category, do not carry a distinct perceptual loudness bias in this listening cohort.

### 3.2 Exploratory Analysis of Individual Consonants

To investigate whether individual consonants exhibit specific loudness contributions that were masked by categorical pooling, data from all 26 stimulus pairings (13 consonants *×* 2 presentation orders) were analyzed individually (ETSB Figure V).

**Figure 5.**
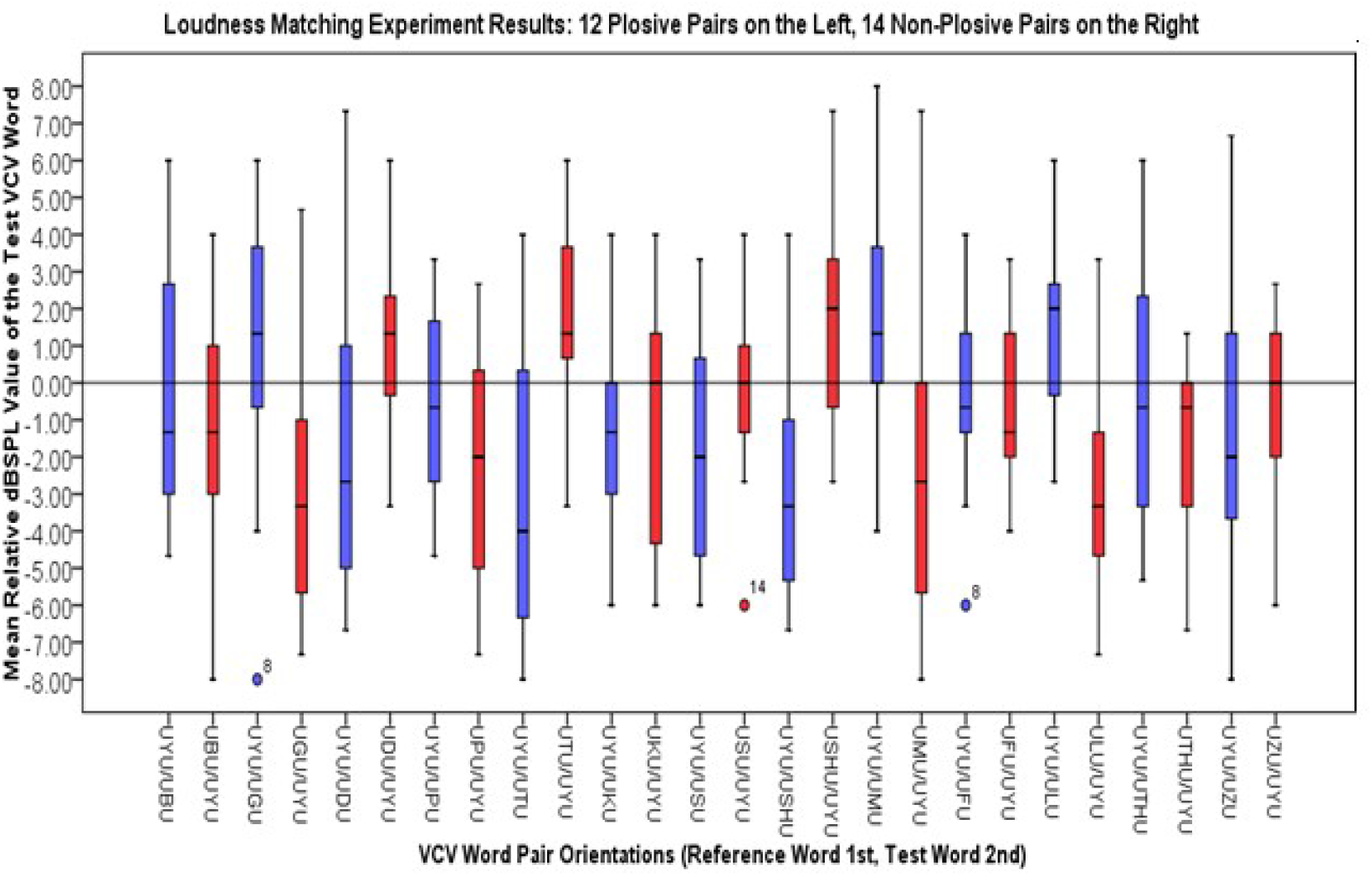
Box and Whisker Plots representing the results from the 26 individual VCV word pair loudness matching experimental arrangements. Blue = VCV word orientation where UYU appeared first. Red = VCV word orientation where UYU appeared second (colour online).

In a two-interval method of adjustment task, a systematic time-order error manifests as a parallel downward shift in adjustment levels across both forward and reverse presentations (i.e., the second interval is consistently perceived as louder, prompting the listener to attenuate whichever sound appears in Interval 2). Conversely, a true stimulus-specific loudness difference relative to /uju/ produces an inverted “mirror pattern”: a stimulus that is inherently louder than /uju/ will be adjusted downward (negative dB offset) when presented in Interval 2, and adjusted upward (positive dB offset) when /uju/ is in Interval 2.

Descriptive evaluation revealed that 7 of the 13 test VCV syllables (/ubu/, /ufu/, /uku/, /upu/, /usu/, /uTu/, /uzu/) exhibited responses dominated by procedural time-order bias. However, 6 VCV syllables demonstrated symmetrical order inversions consistent with distinct consonantal loudness weightings (Table 6):

**Table 6:**
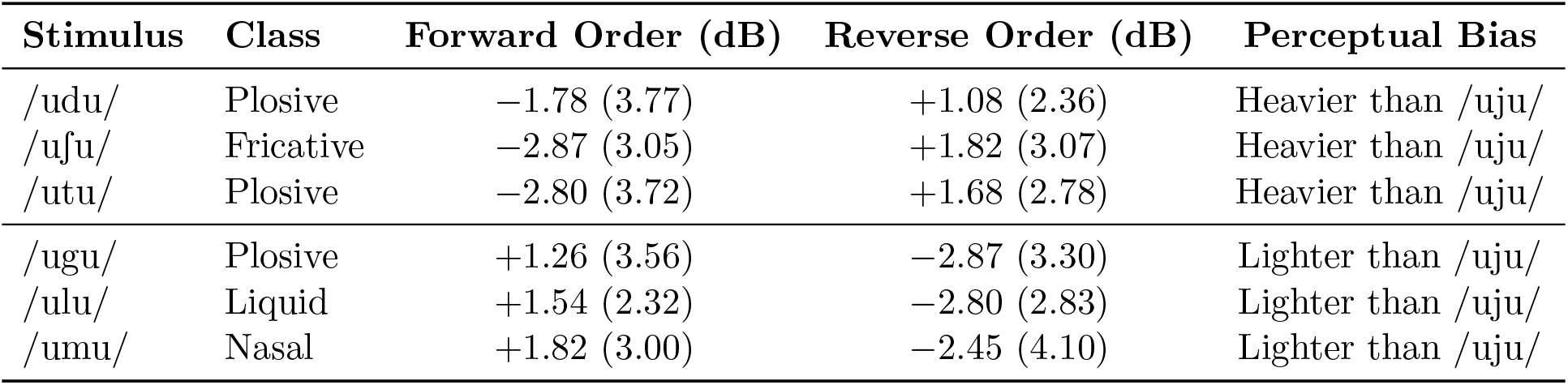
Mean adjustment levels (*±*SD) for the six VCV syllables exhibiting symmetrical order inversion relative to the /uju/ reference.

- **Heavier Perceptual Weighting:** Syllables /udu/, /uSu/, and /utu/ were systematically attenuated in forward presentation (mean offset range: *−*1.78 dB to *−*2.87 dB) and increased in reverse presentation (mean offset range: +1.08 dB to +1.82 dB).
- **Lighter Perceptual Weighting:** Syllables /ugu/, /ulu/, and /umu/ required positive amplification in forward presentation (mean offset range: +1.26 dB to +1.82 dB) and attenuation in reverse presentation (mean offset range: *−*2.45 dB to *−*2.87 dB).

Planned paired *t*-tests comparing forward versus reverse adjustment levels for these two subgroups confirmed significant bidirectional shifts for both the heavier subgroup (*t*(18) = 2.84, *P* = 0.01) and the lighter subgroup (*t*(18) = 4.12, *P <* 0.001). Crucially, these 6 syllables span four distinct phonetic classes (2 voiceless/voiced plosives, 1 voiceless fricative, 1 nasal, 1 liquid), confirming that these psychophysical offsets reflect token-specific acoustic-phonetic characteristics rather than a general plosive advantage.

## 4 Discussion

The present study investigated whether plosive consonants possess an inherent perceptual loudness weighting that exceeds that of non-plosive consonants in symmetrical VCV syllables presented to hearing-impaired listeners. The omnibus statistical analysis demonstrated no significant difference in equal-loudness adjustment levels between plosive and non-plosive syllables (*F* (1, 18) = 0.006, *P* = 0.94). Consequently, the experimental hypothesis is rejected. The finding that plosives do not systematically bias perceived syllable loudness indicates that earlier observations of plosive loudness advantages obtained via subjective rating scales in consonant-vowel (CV) contexts (Sharf, 1971) do not generalize to intervocalic (VCV) loudness balancing tasks in listeners with sensorineural hearing loss.

### 4.1 Categorical versus Token-Specific Loudness Effects

While the categorical comparison across manners of articulation was null, exploratory analysis of individual phonemes revealed that 6 of the 13 test syllables (/d/, /g/, /l/, /m/, /t/, /S/) exhibited consistent bidirectional matching offsets across presentation orders. These offsets corresponded to physical amplitude adjustments of 1 to 3 dB to achieve equal loudness with the /uju/ reference.

Crucially, these 6 consonants do not align with any single phonetic category: they comprise three plosives (/d/, /t/, /g/), one fricative (/S/), one liquid (/l/), and one nasal (/m/). Furthermore, the direction of the perceptual bias differed within the plosive class itself: /t/ and /d/ were adjusted downward (indicating higher perceived loudness relative to the reference), whereas /g/ required positive gain (indicating lower perceived loudness). This within-class divergence directly undermines the hypothesis of a uniform class-wide weighting and suggests that the observed shifts reflect token-specific acoustic properties—such as local spectral peak prominence relative to the listener’s audiometric profile or specific consonant-to-vowel transition acoustics—rather than broad manner-of-articulation effects.

These findings provide nuanced support for models of speech loudness. Historically, speech loudness perception was considered almost entirely vowel-dominated, with consonantal energy exerting negligible influence when vowel levels were fixed (Lehiste and Peterson, 1959; Montgomery et al., 1987). Our data demonstrate that while the normalized flanking vowels established the predominant baseline loudness, manipulating consonant level produced measurable, token-specific offsets in syllable loudness matching, corroborating the findings of Orr et al. (2010). However, temporal integration alone (Moore, 2014) cannot fully explain these offsets; consonant duration (Table 3) did not systematically correlate with the direction or magnitude of the adjustment levels across the 6 significant tokens.

### 4.2 Methodological Considerations and Limitations

Several methodological factors must be considered when interpreting these findings:

- **Time-Order Effects:** A primary challenge in the experimental design was the presence of time-order bias, a well-documented artifact in two-interval method-of-adjustment paradigms (Florentine et al., 2011; Oberfeld, 2015). For 7 of the 13 test syllables, loudness judgments were dominated by order effects (Interval 2 perceived as louder), obscuring potential subtle phonemic loudness differences.
- **Phonetic Context:** Stimuli were restricted exclusively to the symmetrical /u/ context (/u/C/u/). The close back rounded vowel /u/ possesses low first and second formant frequencies, which may create specific masking or contrast effects with adjacent consonants that differ from open or front vowel contexts (e.g., /a/ or /i/).
- **Stimulus Generation and Talker Characteristics:** All stimuli were produced by a single male talker. Because phonetic acoustic cues vary substantially across talker gender, age, and vocal effort (Brandt et al., 1969), the token-specific adjustments observed here cannot be assumed to apply uniformly across all talkers.
- **Listening Cohort:** Participants presented with mild-to-moderate high-frequency sensorineural hearing loss and were tested unaided at a conversational level (65 dB SPL). While this cohort represents typical presbycusis profiles, reduced audibility in the high frequencies likely attenuated the perceived loudness of high-frequency consonantal bursts (e.g., /s/, /k/) relative to lower-frequency voiced transitions.

### 4.3 Implications for Clinical Audiology and Loudness Modeling

From a clinical perspective, these empirical results suggest that broad adjustments to prescriptive hearing aid gain targets or speech audibility models (such as the speech banana) based on consonant manner of articulation are unwarranted. Because the observed loudness differences were token-specific and limited to 1–3 dB, categorical corrections for plosive loudness would risk over- or under-amplifying specific phonemes without improving overall loudness balance. In modern digital hearing aids equipped with multi-channel wide dynamic range compression (WDRC) and fast-acting transient management, syllable loudness is governed dynamically by incoming spectral distributions rather than static phonemic categories.

## 5 Conclusions

In summary, this study tested whether plosive consonants possess an inherent perceptual loudness weighting exceeding that of other consonant classes in listeners with sensorineural hearing loss. In a prospective loudness matching task using symmetrical VCV syllables, no statistically significant difference was observed between plosive and non-plosive syllables as broad categories. Although individual phonemes exhibited modest token-specific loudness offsets of 1 to 3 dB, these did not track manner of articulation. These findings indicate that speech loudness in symmetrical syllables is primarily anchored by vowel energy, with secondary phonemic influences operating at the level of specific acoustic tokens rather than broad phonetic classifications.

